# Quadriceps and hamstrings coactivation in exercises used in prevention and rehabilitation of hamstring strain injury in young soccer players

**DOI:** 10.1101/574210

**Authors:** Gonzalo Torres, David Chorro, Archit Navandar, Javier Rueda, Luís Fernández, Enrique Navarro

## Abstract

This study aimed to study the co-activation of hamstring-quadriceps muscles during submaximal strength exercises without the use of maximum voluntary isometric contraction testing and compare (i) the inter-limb differences in muscle activation, (ii) the intra-muscular group activation pattern, and (iii) the activation during different phases of the exercise. Muscle activation was recorded by surface electromyography of 19 elite male youth players. Participants performed five repetitions of the Bulgarian squat, lunge and the squat with an external load of 10 kg. Electrical activity was recorded for the rectus femoris, vastus medialis, vastus lateralis, biceps femoris and semitendinosus. No significant inter-limb differences were found (F_1, 13_=619; *p*=0.82; partial η^2^=0.045). Significant differences were found in the muscle activation between different muscles within the muscle group (quadriceps and hamstrings) for each of the exercises: Bulgarian squat (F_1,18_=331: *p*<0.001; partial η^2^=0.80), lunge (F_4,72_=114.5; p<0.001; partial η^2^=0.86) and squat (F_1,16_=247.31; *p*<0.001; partial η^2^=0.93).Differences were found between the concentric, isometric and eccentric phases of each of the exercises (F_2, 26_=52.27; *p*=0.02; partial η^2^=0.80). The existence of an activation pattern of each of the muscles in the three proposed exercises could be used for muscle assessment and as a tool for injury recovery.

## Introduction

Performance in soccer depends on psychological, physiological and biomechanical factors (1, 2). The study of these factors not only helps improve performance, but also works towards injury prevention. Performance and injury prevention are not isolated fields, and the presence of an injury can affect on-field performance. Measurement of the muscular capacity of players is an important factor in the evaluation and prediction of the functional capacity of the player (3).

Muscular capacity has been traditionally measured using isokinetic machines (4, 5). However, recently many studies have used electromyography (EMG) (6–8). One of the major aims of such studies has been to to determine asymmetry in soccer players, which can affect both performance (9, 10) and injury (11). Such comparative studies have been carried out in youth soccer players as well (12–15) finding that there may be a neuromuscular pattern and that force work could be related to a reduction in the injury rate in football players.

Asymmetry in soccer players could be offset with proper exercise prescription, which facilitates improvements in musculoskeletal function by addressing the specific needs of the subject as an integral part of any rehabilitative, preventive, or maintenance program (16). Functional weight-bearing exercises have received a significant amount of attention as the preferred mode of exercise for lower extremity strengthening (17). In studies with EMG, basic strength exercises (forward lunge, bugarian squat, lateral step-ups, squat) are frequently used so that they are easily replicable. These are simple exercises used to strengthen the quadriceps and hamstrings, and their study could provide information about the activation of different muscles of the quadriceps and hamstrings and to differentiate the activation differences in the exercise phases (Caterisano, Moss (18)). Considering the muscle group that they target, these exercises can be crucial for soccer players given that recent research has shown that monitoring the electrical activity of individual muscles could help in the injury rehabilitation and prevention process (18, 19).

Specifically, in the case of soccer players, quadriceps and hamstrings account for 19% and 16% of all injuries respectively (20). The case of hamstring strain injury is of concern given that there is an increase of 4% every year in the male teams (21). Muscular imbalances have been cited as an important risk factor for hamstring strain injury (22, 23). This has been traditionally assessed by evaluating muscular imbalances through isokinetic machines (24–26), although recent studies have begun using surface electromyography (EMG) to evaluate muscular imbalances in different strength exercises. EMG has been used to evaluate the nordic hamstring curl (27, 28), the leg curl (29, 30), bilateral open chain (31) and unilateral closed chain exercises like forward lunge, bulgarian squat and unilateral bridge (29, 32). Research has shown, these exercises targeting the quadriceps and hamstrings have proved beneficial not only for these muscle groups, but also in rehabilitation and prevention of anterior cruciate ligament (33) patellofemoral pain syndrome (34) or groin injuries (35). However, these researches have used adult/amateur athletes, but none have specifically focused on young soccer players, which have traditionally used isokinetic machines(12).

Studies using EMG have required the normalization of EMG signal (36). Many normalization techniques are available being the maximal voluntary isometric contraction (MVIC) the most used standard methods (36). However some controversy exists about its reliability (37–39) leading researchers to study the agonistic-antoagonist coactivation of muscles as an alternative by normalizing the electrical activity of one muscle in relation with the other (18). This can be useful because it gives information on how muscles work synergistically and permit a normalization that is not demanding for the athlete, facilitating its use at any time of the season.

Hence, this study aimed to study the hamstring-quadriceps muscle co-activation during submaximal strength exercises (Bulgarian squat, forward lunge and squat) without the use of voluntary maximum isometric contraction testing and compare (i) inter-limb differences in muscle activation, (ii) intra-muscular group activation pattern, and (iii) activation during different phases of the exercise. The following hypotheses were raised:

1. Inter-limb differences would be noted for the different exercises.
2. Muscle activation is different according to the phases of movement.
3. There is a different pattern of intra-hamstrings and intra-quadriceps muscle group co-activation for each exercise.

## Material and methods

### Participants

Nineteen soccer players participated in the study (Table 1), all of them men, belonging to the youth soccer team of a professional Spanish team.

**Table 1.**
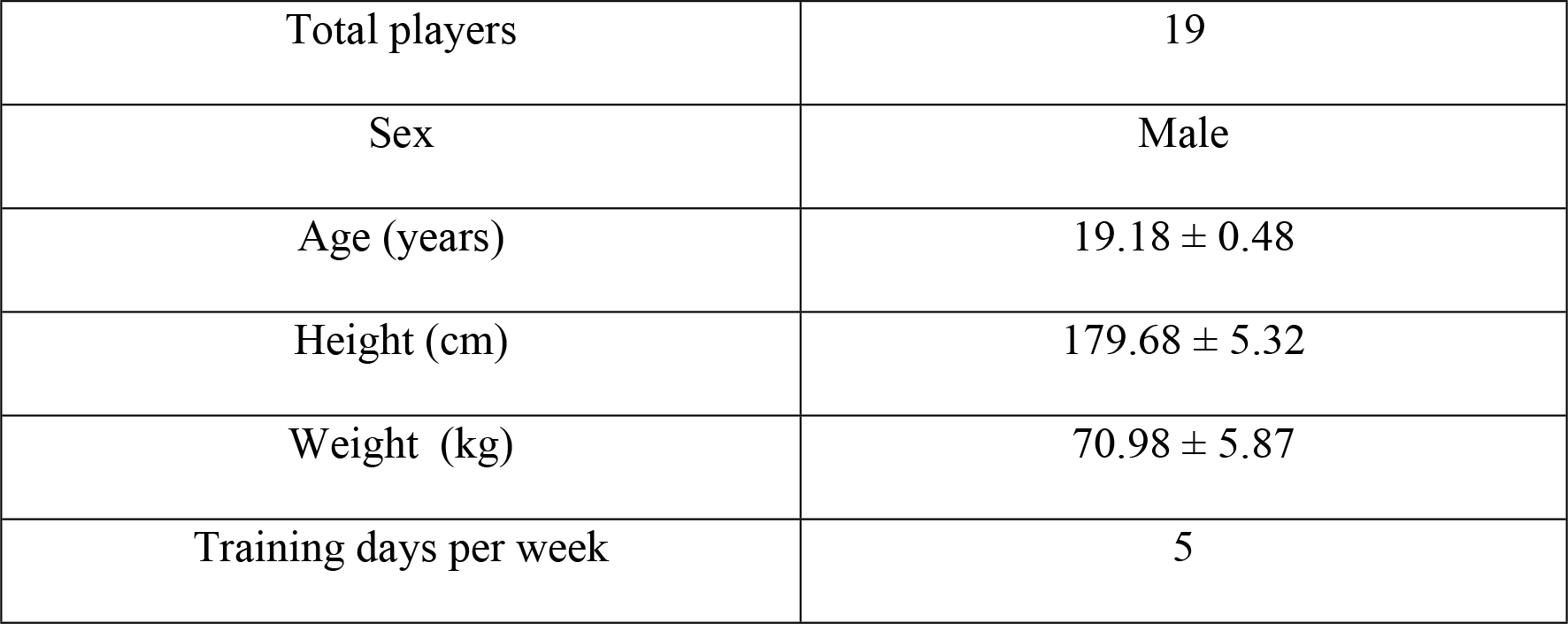
Basic demographic and descriptive data of the included sample (Mean ±SD)

The players who were included in the study did not sustain leg injury in the last six months prior to the day of testing. All participants gave their written informed consent to participate in the study. The study followed the guidelines of the Declaration of Helsinki and has been approved by the ethics committee of the **blinded for peer review**.

### Study Design

A descriptive study was carried out in a sports biomechanics laboratory, where participants were required to perform the Bulgarian Squat, Squat and Lunge in a single session.

1. 1. Lunge. The initial starting position was with one forward leg, straight trunk and arms to the side of the body. The lunge exercise consisted in advancing one leg in front of the other, and then flexing both knees without touching the ground while the trunk was straight. The forward knee reached a flexion angle of 90°. It was ensured that the knee of the advanced leg did not surpass the toe of the foot (6).
2. 2. Bulgarian squat. The initial starting position consisted in advancing one leg while the back leg was placed on an elevated surface of 50cm. The players then had to flex the forward knee while keeping the trunk straight (40).
3. 3. Squat. The participant’s feet were separated such that they were apart at shoulders’ length, and the players had to lower themselves until they reached a 90° knee flexion angle while the trunk was straight (41, 42).

### Procedure

The players first warmed up under the guidance of a strength and conditioning coach of the club. It consisted of seven minutes of running followed by joint mobility and core activation exercises for an additional three minutes. Finally the players familiarized themselves with the three exercises at the end of the warm-up (43).

After warming-up, EMG sensors were placed on the rectus femoris (RF), vastus medialis (VM), vastus lateralis (VL), bíceps femoris (BF) and semitendinosus (ST) with the players’ skin being shaved and cleaned with alcohol prior to sensor placement. The Seniam protocol was used for sensor placement (44) and recommendations of Ramírez and Garzón (45) were followed, with the orientation for the rectus femoris being 0°, 70° for the vastus medialis and −20° for the vastus lateralis, taking as reference a line drawn from the origin and to the insertion.

EMG data was recorded using Trigno™ Wireless System (Delsys, Inc. Massachusetts, U.S.A). This system allowed players to perform the exercises with complete freedom of movement and was composed of a receiver device, a data registering software, and a series of surface wireless electrodes placed in each of the muscles, plus one placed on the back to be used as an accelerometer to discern the different phases of the exercises. The sensors measured 0.037m × 0.026m × 0.015m, with a distance between electrodes of 0.01m and weighed 0.014 kg and could record both the EMG signal and triaxial accelerometry, with an autonomy of 8 hours and a charging time of 2 hours. The input range was 0.011V, 16-bit resolution, bandwidth between 20-450Hz. Data was captured at 1500Hz. The signal gain of the electrodes was 909V/V±5%. The EMGWorks Acquisition software (Delsys, Inc. Massachusetts, U.S.A) was used to visualize and record the data.

Once the sensors were placed, the players performed the three exercises. Each exercise had three well-differentiated phases, the first phase of descent, the second one isometric phase and finally the phase of ascent. Each exercise was repeated five times (for each leg) and the execution rhythm was externally marked by a stopwatch: two seconds of descent, two seconds of isometric and two seconds of ascent. The external load was 10kg (two 5 kg dumbbells). Between each exercise there was a complete rest of 2.5 minutes.

### Data Processing

The EMGWORKS^®^ software (Delsys, Inc., Massachusetts; U.S.A.) was used to process the data. The first step of the data treatment was the filtering of the signal using a 2^nd^ order, bandpass Butterworth filter (36) with an attenuation of 40dB and a frequency cut between 20-30 Hz (46). A Root Mean Square (RMS) (47) with a window width of 0.05s and a window overlap of 0.025s was later applied to the filtered signal and the signal offset was removed.

To identify the phases of the exercises, the accelerometer signal from a sensor placed on the back of the players in the longitudinal axis was used (48). The EMG and accelerometer signals were superimposed, and the mean RMS of the 5 repetitions was used for each of 3 exercises for analysis.

### Statistical Analysis

Four independent variables were defined: preferred limb, muscle group, phases of the exercise, and exercise and two dependent variables: intragroup muscular ratio and electrical activation (V) were compared. The intragroup muscular ratio expresses the activation of each muscle with respect to the total surface muscular activity of each muscle. For example, the intragroup muscular activation for the RF = Muscular activity of RF/(Muscular ativities of RF+VL+VM).

The statistical analyses were carried out with SPSS 23.0 (Spss Inchicago, IL, USA). An analysis of variance was carried out study the main effects of the 4 independent variables (leg (2) × muscle (5) × phases (3) × exercise (3)). Subsequent post-hoc comparisons were done with Bonferroni corrections. The significance level was set at 0.05 and effect sizes were determined using partial eta squared values (threshold values: small= 0.2, medium = 0.6 and large = 0.8, (49)).

## Results

### Inter-limb effect

Both the main effect of the leg factor (F_1, 13_=619; *p*=0.82; partial η^2^=0.045) and the differences in intra-muscular group activation between dominant and non-dominant leg in each phase of the movement of each muscle in the exercises analysed were not significant. Therefore, for the successive comparisons the mean activation between the two legs was used as the dependent variable.

### Intra-muscular group activation pattern

Significant differences were found in the muscle activation between different muscles within the muscle group (quadriceps and hamstrings) for each of the exercises (Table 2): Bulgarian squat (F_1,18_=331; *p*<0.001; partial η^2^=0.80), lunge (F_4,72_=114.5; p<0.001; partial η^2^=0.86) and squat (F_1,16_=247.31; *p*<0.001; partial η^2^=0.93). In the quadriceps, the post-hoc comparisons showed that the vastus medialis showed the highest activity, followed by the vastus lateralis and then the rectus femoris for all exercises (*p*<0.001). In the hamstring muscles, the semitendinosus had greater activation than the bíceps femoris (*p*<0.001) for the Bulgarian squat and lunge exercises. However, in the squat no significant differences were obtained between the two muscles (*p*=0.175). Comparisons between each muscle in the different exercises showed significant differences in the activation pattern (F_8,128_=2; *p*=0.03; partial η^2^=0.93) (Fig 1).

**Table 2.**
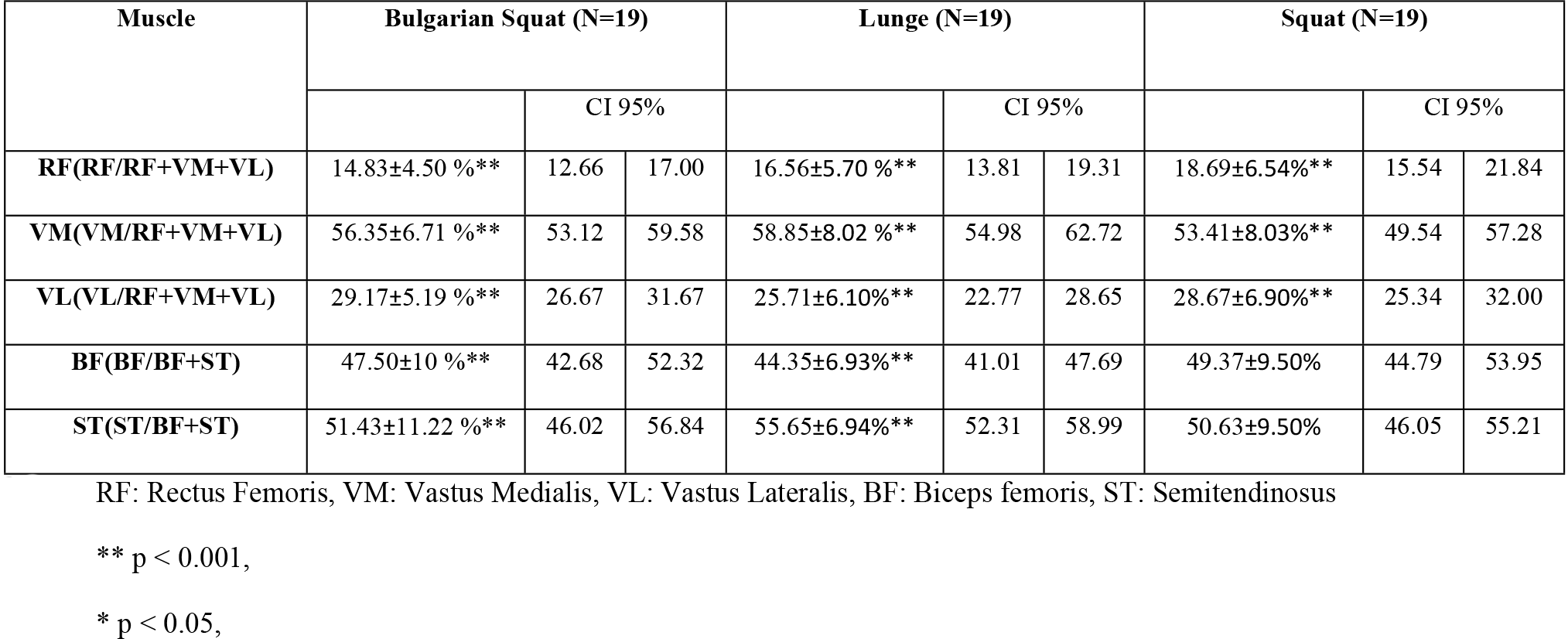
Intra-muscular group activation (Mean± ST) in percentage.

**Fig 1.**
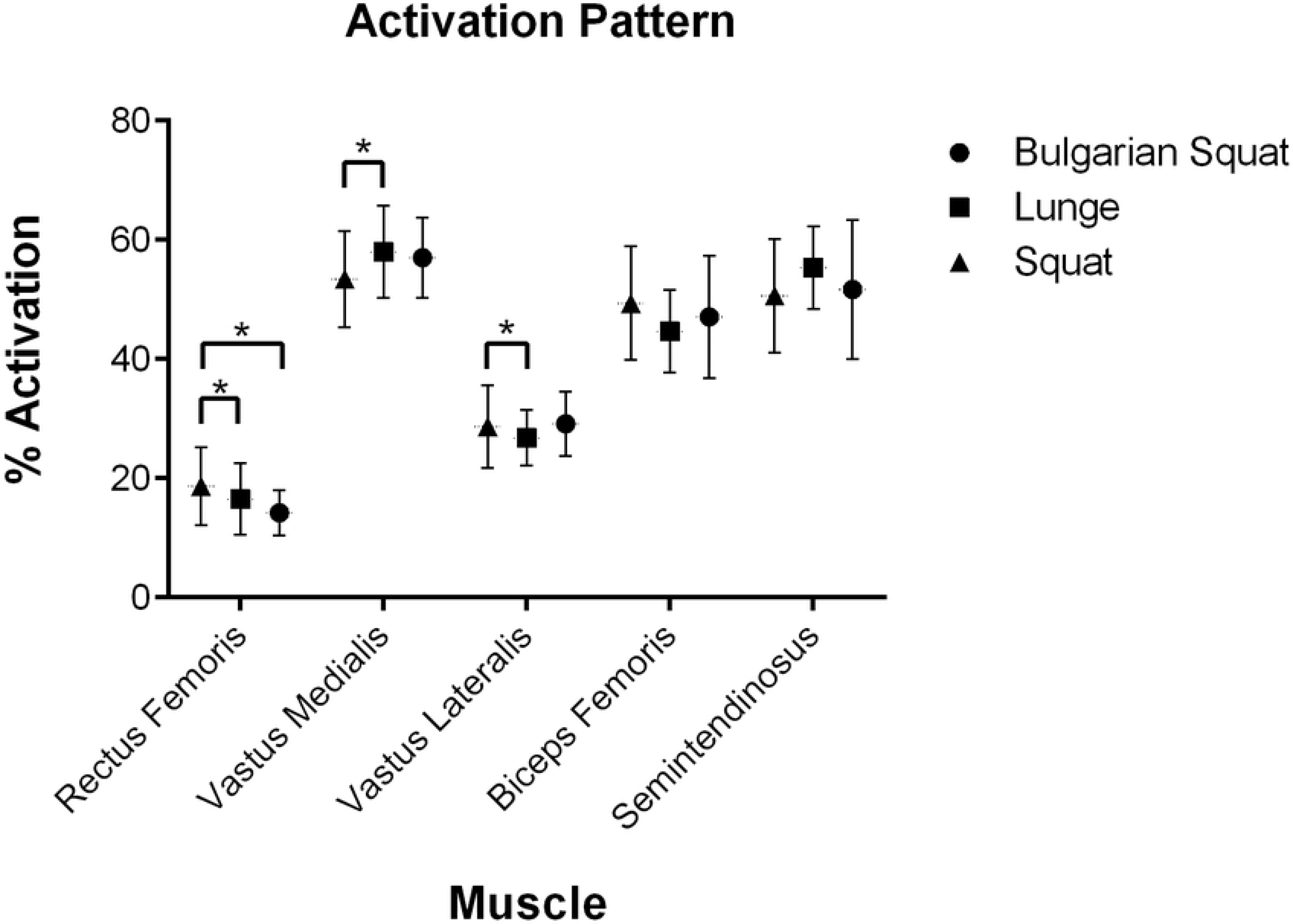
Intra-muscular group activation (mean± standard deviation) in percentage.

### Electrical activation during the movement phases

Significant differences were found in the electrical activity (RMS) between the phases and in the three exercises (F_2, 26_=52.27; *p*=0.02; partial η^2^=0.80).

In the Bulgarian squat exercise (Fig 2), significant differences were found between phases (F_10,58_=12.18; *p*<0.001; partial η^2^=0.68). In all quadriceps muscles, the isometric phase was the one that registered the greatest electrical activity (*p*=0.01). The ascent phase had greater activation than the descent phase (*p*=0.03). In the hamstrings, the ascent phase was greater than the other two (*p*=0.02) in both biceps femoris and semitendinosus.

**Fig 2.**
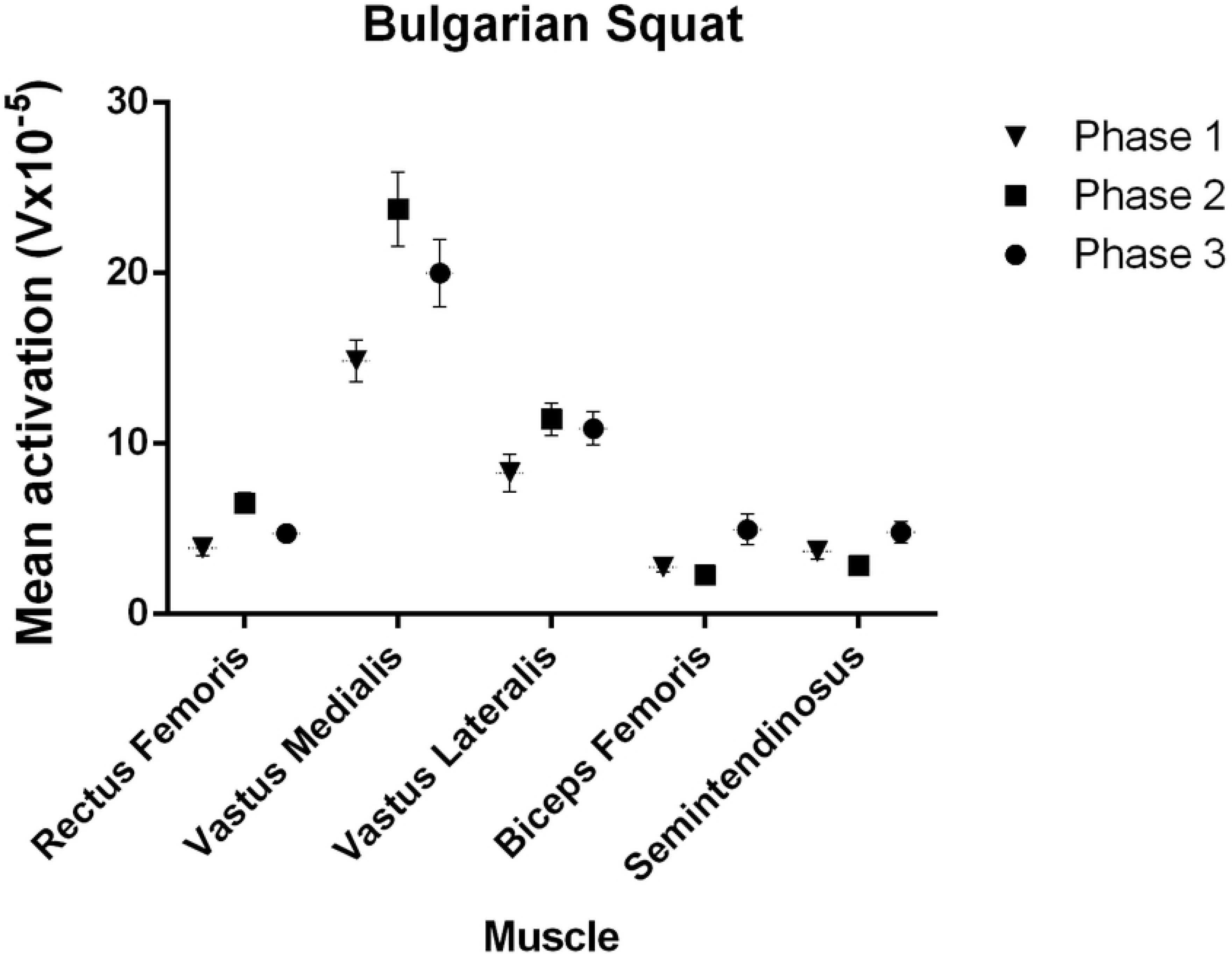
Mean activation (V×10^−5^) of all the muscles in each phase in Bulgarian squat.

In the lunge exercise (Fig 3), significant differences were found between the 3 phases (F_10,54_=16.85; *p*<0.001; partial η^2^=0.76). In the rectus femoris, vastus medialis and vastus lateralis, the isometric phase had greater activation than the descent and ascent phase (*p*<0.001), the ascent phase had greater activation than the descent phase (*p*<0.001). In the biceps femoris, the ascent phase had greater activation than the isometric phase and the descent phase (*p*<0.001); the descent phase had greater activation than the isometric phase (*p*<0.001). In the semitendinosus, descent and ascent phase had similar activation (*p*=0.07).

**Fig 3.**
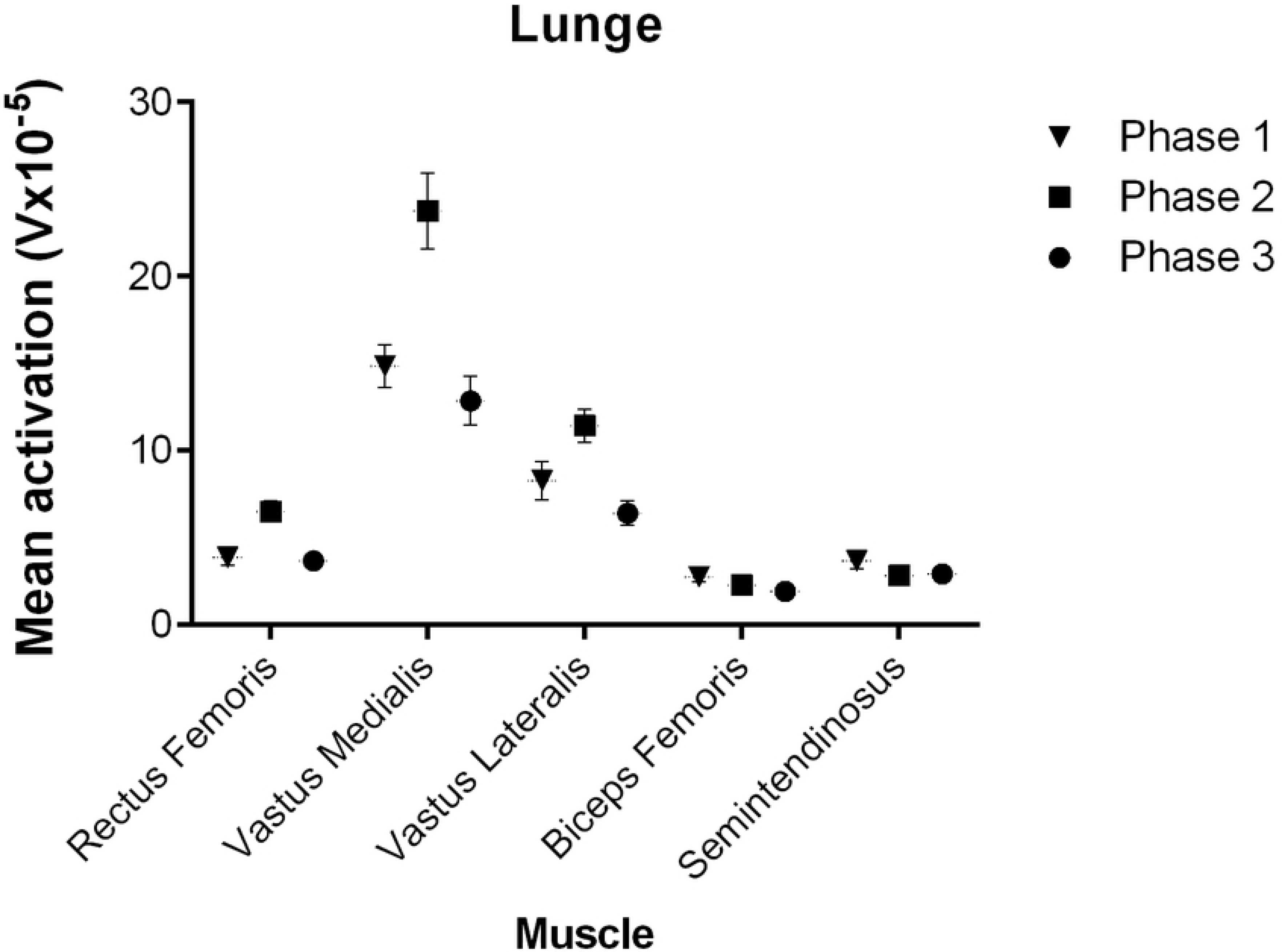
Mean activation (V×10^−5^) of all the muscles in each phase in lunge.

In the squat exercise (Fig 4) there were significant differences between the phases (F_10,46_=8.04; *p*<0.001; partial η^2^=0.64). In the rectus femoris, the isometric phase had greater activation than the descent and ascent phase (*p*=0.04), there were no significant differences between ascent phase and descent phase (*p*=0.09). In the vastus medialis the isometric phase had greater activation than the descent phase (*p*<0.001), there were no significant differences between the isometric phase and the ascent phase (*p*=0.25), the ascent phase had greater activation than the descent phase (*p*=0.04). In the vastus lateralis, the isometric phase had greater activation than the descent phase (*p*= 0.02), the ascent phase had greater activation than the descent phase (*p*=0.03), there was no difference between the isometric and ascent phase (*p*=0.87). In biceps femoris and in the semitendinosus, the ascent phase had the highest activation (*p*<0.001).

**Fig 4.**
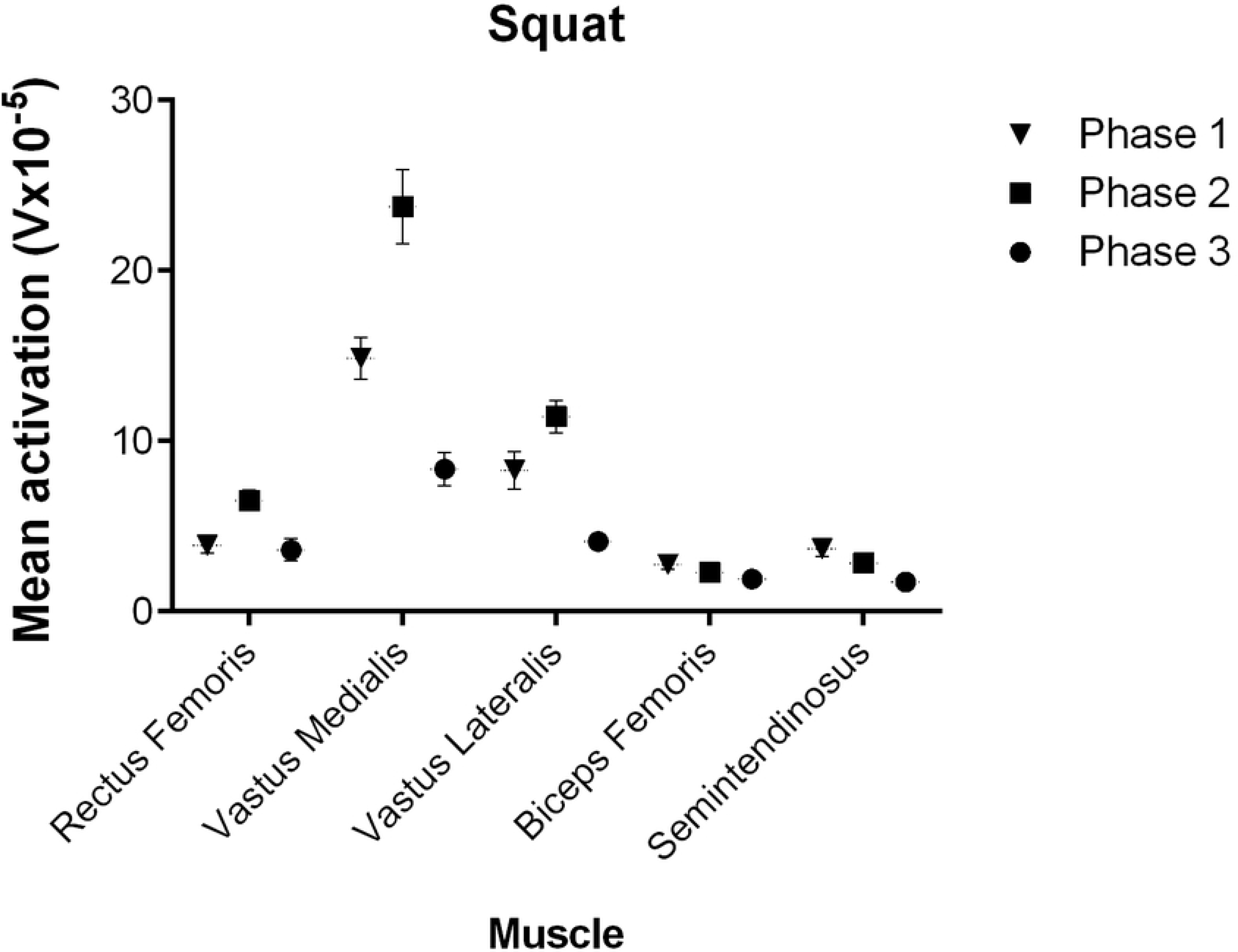
Mean activation (V×10^−5^) of all the muscles in each phase in squat.

### Discussion and implications

In this paper, the surface muscular activation of two important muscular groups, hamstrings and quadriceps, was measured in strength exercises in elite youth soccer players. These exercises, namely the Bulgarian squat, the lunge and the squat, are frequently used and easily replicable. No significant inter-limb differences were observed, rejecting hypothesis 1. Muscle activation patterns showed differences across the intra-muscular group to which they belonged, confirming hypothesis 2. Also, muscle activation differed across the concentric, isometric and eccentric phases of each exercise, confirming hypothesis 3. This is one of the first investigations to study the muscle activation pattern in elite youth soccer players and has important applications where differences in the muscle activation pattern could indicate muscular deficiencies leading to other muscles assuming part of that deficiency.

### Inter-limb effect

The results show that no significant differences were found between dominant and non-dominant leg in electrical activity, other previous studies obtained similar results with professional or semi-professional samples (50–52). Although soccer is an asymmetric sport, the modern demands of the sport lay an emphasis on training both limbs equally. Another important factor to consider is that the participants in this study were elite youth players, playing in national and European competition, and studies have suggested that inter-limb differences in strength decrease with an increase in level of play (53). Inter-limb differences could be a precursor to injuries (54) hence the monitoring of inter limb differences could be important in injury prevention programs.

### Intra-muscular group activation

In all exercises, the muscular activation patterns of the quadriceps muscles were repeated, with the vastus medialis being predominant, followed by the vastus lateralis and finally the rectus femoris (Fig 1). These results were similar to those obtained with amateur athletes previously for the lunge (55) and squat exercises (18, 56). The similar pattern of activation of quadriceps and hamstrings could indicate a synergy of work in the proposed exercises (57). The results showed that there is a very similar activation pattern in the three exercises and in all the muscles (Fig 1), this pattern can be used for the rehabilitation process of an injury or to find deficiencies in the coactivation inside each muscle. This can be applied in the case of the patellofemoral pain syndrome, where vastus medialis activation is the primary goal in the early stages of recovery (58).

In the hamstrings, the greater intra-muscular activation (Fig 1) of the semitendinosus muscle compared to that of the biceps femoris account for its predominance (59, 60). A balance in the synergy of the hamstring muscles is important in the reduction of risk of a potential injury (60). This synergy helps to reduce the tension on the biceps femoris (19, 61), which plays a fundamental role in the hamstring injury since biceps femoris is affected in 80% of all hamstring lesions (62). A low hamstring to quadriceps ratio is believed to be a risk factor for injuries (63) and many rehabilitation and prevention programs focus on strengthening the hamstrings. The heterogeneity of activation pattern of different exercises must be taken into consideration when developing hamstring strength (30), and these exercises help in the coactivation of the hamstring and quadriceps muscle groups.

### Activation during the movement phases

Differences were found between the different phases of the 3 exercises (Fig 2–4), for quadriceps muscles, the greatest activation was recorded in the isometric phase, while in the hamstrings the greatest activation was in the ascent phase, or knee extension phase.

In the quadriceps, in all the muscles and exercises (Fig 2–4) greater activation was found in the ascent phase (concentric) than in the descent phase (eccentric), coinciding with previous studies (64–66). In the concentric phase, there is a greater activation generated as there is a greater recruitment of fibres (67); while in the eccentric phase, a greater proportion of rapid contraction motor units were active at the expense of less recruitment of other motor units for load reduction (68). This finding illustrates the contribution of the hamstrings to effective force production in the concentric phase of the squat. There are two major theories to justify less activation in the eccentric phase than in the concentric phase. It is believed that the lower amplitude of EMG during eccentric phases may be due to the greater predominance of muscle-tendon strength produced by elongation in the eccentric phase (69). Other authors, however, have proposed that differences in EMG amplitude may be due to different neuronal pathways for each activation (68).

The isometric phase showed that rectus femoris presented the least activation in the quadriceps in all exercises, giving predominance in the stabilization of movement to the vastus medialis and vastus lateralis (67).

For hamstrings muscles, there was greater activation in all phases in the Bulgarian squat (Fig 2) than in the normal squat (Fig 4), which may be due to a greater demand for co-activation of the posterior leg musculature to keep joints and the body stable (70). Wright, Delong (60) emphasized the importance of the posterior muscles in exercises such as the Bulgarian squat, where the knee joint was fixed, thereby increasing the activation of the hamstrings.

These results can help sports scientists and strength and conditioining coaches, especially those working with youth players, when they want to influence a greater muscle activation for hypertrophy or implement these exercises in processes of strength development and rehabilitation.

### Choice of exercises

The proposed exercises have been considered suitable for muscle evaluation and rehabilitation (6, 71, 72) and have therefore been implemented in most soccer club training programmes by physical trainers. The in-depth study of such exercises with a specific high-performance sample may allow soccer clubs to implement specific protocols for the development of lower train strength in players. An ineffective recovery of the thigh injury could cause a relapse in the soccer (73), so it is vital to be able to correctly quantify the recovery process where EMG can be used as a reliable method. Hence, the monitoring of such exercises could be used by sports scientists and strength and conditioning coaches. Specific leg exercises can be graded by exercise intensity providing atheltes and therapists to select appropriate exercises during different phases of prevention.

## Conclusion

EMG activation patterns in different muscles differed across the muscle group to which they belonged. Comparing the different phases of the exercises, in the quadriceps the isometric phase had the greatest activation followed by the concentric and eccentric phases; while in the hamstrings the concentric phase was the one with the greatest activation. A monitoring of the 3 proposed exercises, using EMG, can be used to find activity deficits in the muscles or for assessing muscles activation during injury rehabilitation. This procedure could be used in the future as a tool for researching about the risk of hamstrings strain injury in soccer.

## Acknowledgments

The authors thank all the experts and medical services for their collaboration in the project. Furthermore, we want to highlight the good involvement of all players in the study.

